# Morphology, biology and phylogeny of *Xenos gadagkari* sp.nov. (Strepsiptera: Xenidae): an endoparasite of *Polistes wattii* (Hymenoptera: Vespidae)

**DOI:** 10.1101/2024.06.18.599476

**Authors:** Deepak Nain, Anjali Rana, Rhitoban Raychoudhury, Ruchira Sen

## Abstract

We report the morphology, biology and phylogeny of a new strepsipteran endoparasitic species: *Xenos gadagkari,* from the primitively eusocial wasp *Polistes wattii* from Punjab, India. We report a high abundance of the endoparasite in *P. wattii* population of our study area and provide the morphological description of adult males, male puparium and neotenic females. This species is sufficiently different from other *Xenos sp* reported from various social wasps and therefore, constitutes a new species, *X. gadagkari* Sen and Nain, sp. nov. Although there are a few reports of Strepsipteran parasites from India, this is the first confirmational report with a thorough description of a *Xenos* species from the Indian population of *P. wattii* and underscores the need for further investigation of the diversity and distribution of this group in India. Our findings contribute to the growing body of knowledge on *Xenos* and provide a foundation for future behavioural, microbial and molecular studies on this enigmatic group of insects.

## Introduction

Strepsipterans are an obligatory endoparasitic group of insects infecting a wide range of other insect species (Cook, 2014; Kathirithamby, 1989, 2009, 2021). The strepsipteran genus *Xenos* has been reported from various species of eusocial wasp genera *Polistes, Mischocyttarus, Belanogaster, Polybia, Apoica, Vespa and Vespula* (Benda *et al*., 2022; Cook, 2019; Dong *et al*., 2022; Kathirithamby, 2024).. *Xenos* infections bring about striking changes in the morpho-physiology and the behaviour of these eusocial wasps (Beani et al., 2011). Generally, a healthy eusocial female wasp can either gain direct fitness by laying eggs or can gain indirect fitness by becoming a worker and helping egg-laying relatives. However, *Xenos* infections prevent both these mechanisms and restrict the host from gaining any such fitness. Infected wasps show marked inhibition of their ovarian development, preventing them from laying eggs (Hughes et al., 2004). These infections also modify the host’s behaviour whereby infected females desert their natal nests and join a nest-free aggregation of other similarly infected conspecifics and subsequently act as an agent to spread the parasite onto the healthy nests. Such aggregations were first observed by W.D. Hamilton (Hughes, 2002) and later studied and reviewed by others (Beani, 2006b; Beani & Massolo, 2007; Cappa et al., 2014; Dapporto et al., 2007; Geffre et al., 2017; Hughes et al., 2003, 2004; Hughes et al., 2004; Kathirithamby, 2009). These behaviours render the wasps unable to carry out the normal worker-like behaviours of rearing the brood of their siblings and thus also prevent the workers from gaining indirect fitness.

To date 46 species of *Xenos* have been reported worldwide with 23 species obtained from *Polistes* (Dong *et al* 2022, World Strepsiptera Database). These include 18 species from South America, 8 species from North America, 3 species from Europe, 6 species from Africa and 11 species from Asia. However, detailed morphological descriptions have not been reported for some of these species (Benda *et al*., 2022; Cook, 2019; Dong *et al*., 2022). Of the many strepsipteran species that have been reported from Asia, *Xenos* has been reported only from three *Polistes* species of the continent - *X. circularis* in *P. rothneyi, X. yamaneorum* in *P. gigas* (Kifune & Maeta, 1985) and *X. hebraei* in *P. olivaceous* (Kinzelbach, 1978). A comprehensive description of the morphological, physiological, and behavioural aspects is also lacking for any *Polistes-Xenos* association from India. *P. wattii* is one of the most abundantly distributed wasps from north India and is found in at least 13 other countries of Asia (Nain & Sen, 2023; Sen et al., 2022). Here we describe the morphological features, biological note and phylogeny of *X. gadagkari* sp. nov., associated with *P. wattii*.

Benda *et al*, (Benda *et al*., 2022) reported the presence of *X. hebraei* in *P. wattii* from Oman assuming that it is the same *Xenos* species reported from *P. olivaceous* (Kinzelbach, 1978) as both species of *Polistes* belong to the same subgenera. However, they did not provide any morphological description or phylogenetic information of the *Xenos sp* found in Oman. A description of male *X. hebraei* was also lacking in the original report by Kinzelbach (1978). We provide the differences in female cephalothorax and morphometry between *X. hebraei* and *X. gadagkari* and show that it is logical to describe *X. gadagkari* as a new species instead of assuming it to be the same as *X. hebraei* found in *P. olivaceous*.

*Etymology: Xenos gadagkari* is named after Prof. Raghavendra Gadagkar, the pioneer of behavioural ecology and wasp biology in India.

## Material and Methods

### Taxonomic identification

6 nests of late colony phase were collected in 2021 (Table 1) from the Indian Institute of Science Education and Research (IISER), Mohali, campus (30.7046° N, 76.7179° E) to check the parasitic occurrence in them. *Xenos* infections were found in both male and female wasps.

**Table 1.** Numbers of stylopized wasps, compared to healthy wasps, in nests at late the colony phase. All nests were collected from IISER Mohali Campus.

| Nest No | Date of Collection | Healthy Male<br>(No externally visible parasite) | Stylopized Male | Healthy Female<br>(No externally visible parasite) | Stylopized Female |
| --- | --- | --- | --- | --- | --- |
| 1 | 27-09-2021 | 46 | 8 | 32 | 0 |
| 2 | 24-11-2021 | 7 | 2 | 12 | 1 |
| 3 | 24-11-2021 | 1 | 0 | 35 | 3 |
| 4 | 19-12-2021 | 5 | 0 | 107 | 3 |
| 5 | 09-10-2022 | 8 | 1 | 34 | 4 |
| 6 | 18-10-2022 | 19 | 6 | 46 | 16 |

For morphological studies, stylopised wasps were collected from nest-free aggregations of parasitized *P. wattii* during May 2022, from Panjab University campus, Chandigarh, India (30.7606° N, 76.7653° E). Seven individuals, from both sexes of *Xenos,* were used for morphological measurements. Infected *P. wattii*, with *Xenos* males, were dissected to yield live male parasites from their puparium for studying the morphological features. A Leica M205C microscope was used for the dissection and measurement of the wasps and their *Xenos* parasites. Measurements were taken from mature (fully sclerotised) males dissected out of puparium. To ensure that the females had reached the maximum possible size, only the females with at least one planidium larva (to ensure the presence of developed reproductive organs and mated status) were selected for measurement. To study the morphological features in greater detail, Scanning Electron Microscopy (SEM) images of the head of a male parasite were taken using a JEOL JSM-6010 PLUS/LV Scanning Electron Microscope.

### DNA Extraction and Phylogenetic tree construction

To identify the phylogenetic position of *X. gadagkari,* molecular phylogenetic analysis was conducted using the partial sequence of the mitochondrial Cytochrome Oxidase I (*CO1*) gene. Three male and three female *Xenos,* obtained from *P. wattii* wasps collected from IISER Mohali campus were used for DNA extraction. DNA was extracted by the Phenol-Chloroform-Isoamyalcohol (PCI) method after crushing the samples in 200µl lysis buffer containing 10mM each of Tris-HCL (pH 8.0), EDTA (pH 8.0) and NaCl. DNA was precipitated using isopropanol, dissolved in 1X TE (pH 8.0) and quantified using the NanoDrop^TM^ 2000 spectrophotometer (Thermo Fisher Scientific). A ∼600bp long fragment of the *CO1* gene was amplified and sequenced using the primer pair CO122F/669R (Benda *et al*., 2021). Polymerase chain reactions (PCR) were performed under the following conditions: an initial denaturation step at 94°C for 5 minutes, 35 cycles of denaturation (94°C, 30 sec), annealing (50°C, 45 sec), extension (72°C, 1 min) and a final extension at 72°C for 5 minutes. PCR products were electrophoresed and visualized on 1% agarose gels and then were cleaned using Exonuclease I and Shrimp alkaline Phosphatase (New England Biolabs Inc.). Both forward and reverse strands were sequenced using BigDye® Terminator v3.1 cycle sequencing kit. Generated sequences have been submitted to NCBI with accession number OR086007-OR086008, PP869407-PP869410).

*X. gadagkari* mitochondrial *CO1* gene sequences were aligned with other known sequences from NCBI using ClustalW multiple alignment tool (Thompson *et al*., 1994) and manually edited in BioEdit 7.2.5 version (Hall, 1999). In all, *CO1* sequences from 30 different species of *Xenidae* were collected from NCBI along with *Stylops advarians* as the outgroup taxon. For phylogenetic tree construction, the best evolutionary model was evaluated in MEGA7 (Kumar *et al*., 2016). Bayesian phylogenetic tree was constructed using GTR+g+i model (general time reversible model with γ-distributed rate variation and a proportion of invariable sites) in MrBayes v3.2.5 (Ronquist *et al*., 2012). Simulations were run until the standard deviation of the split frequency came below 0.01. Resulting phylogenetic trees were visualized in Figtree v1.4.2 (Rambaut, 2009). The sequence divergence between *X. gadagkari* and *X. vesparum* was calculated in DnaSP v6 (Rozas et al 2017).

## Results

### Biological Note

*P. wattii* follows a biannual nest founding strategy. The overwintered wasps initiate solitary foundress nests in spring (March) but these are abandoned within 2-3 months. Nests are again initiated in summer by multiple females and last till October-November (Sen *et al*., 2022). *Xenos* infection was spotted in wasps throughout their social cycle (March-December) as well as in hibernating wasps (December-February) (Nain et al unpublished data). Nest-free aggregations of stylopised wasps were found late April-May. In April 2015, a small cluster (10 wasps) of parasitized wasps were found in IISER-Mohali campus. It has been proposed that *X. vesparum-*infected *P. dominula* females cluster to facilitate the mating of the parasite (Hughes *et al*., 2004). Congruent with that hypothesis, both male (male pupa or empty puparium) and female *Xenos* were found in the aggregating wasps. As is typical in the case of most other strepsipterans, adult *Xenos* males eclose and leave the host while the neotenic females remain inside the host. Various developmental stages of the parasite were found in the wasps. Infected adult wasps with single or multiple parasites (Fig 1-2) were found in the nests during the late colony phase, in Oct-Nov (Table 1).

**Figure 1.**
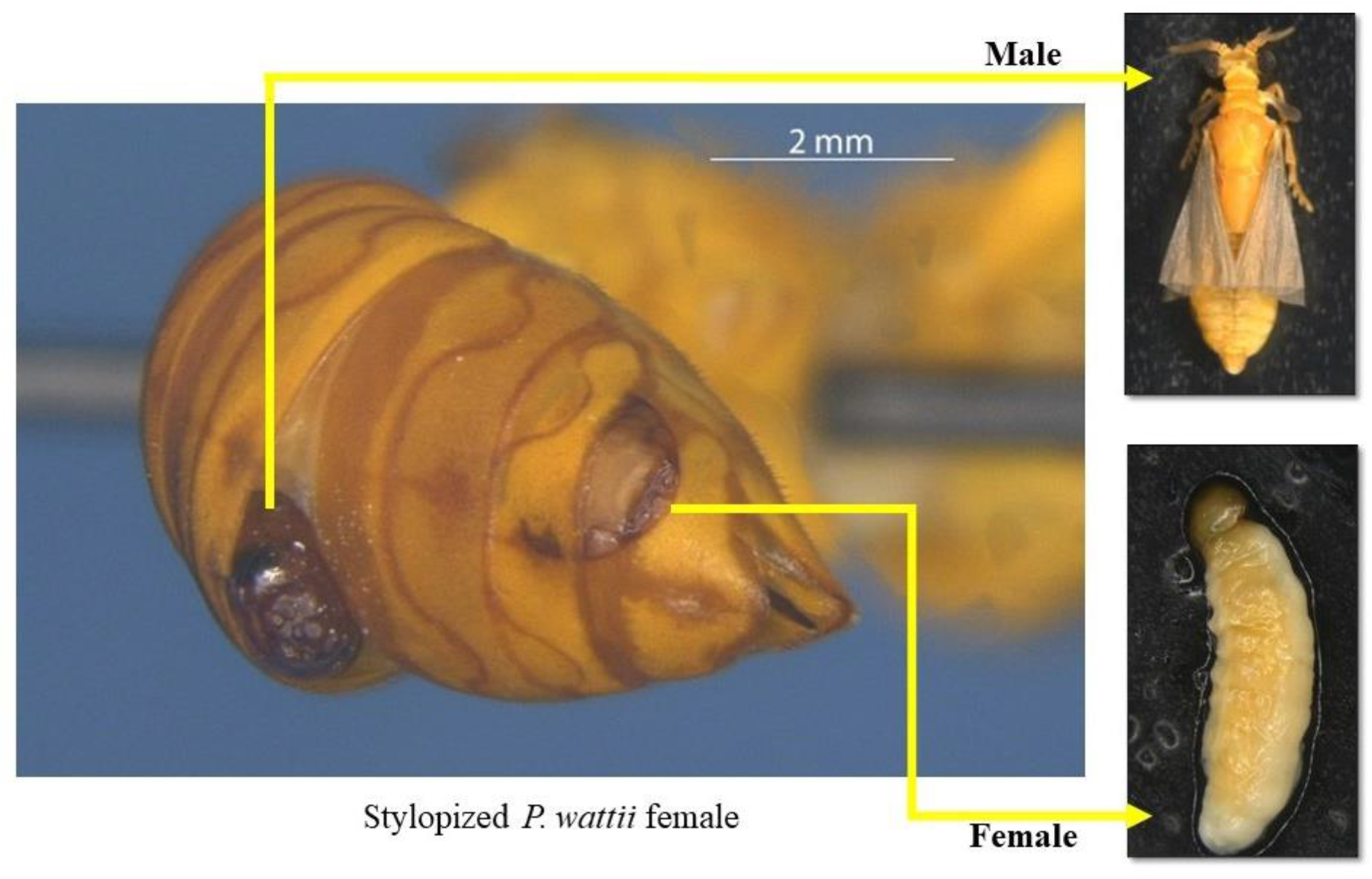
Abdomen of a stylopized *P. wattii*, infected with a male and a female *X. gadagkari*.

**Figure 2.**
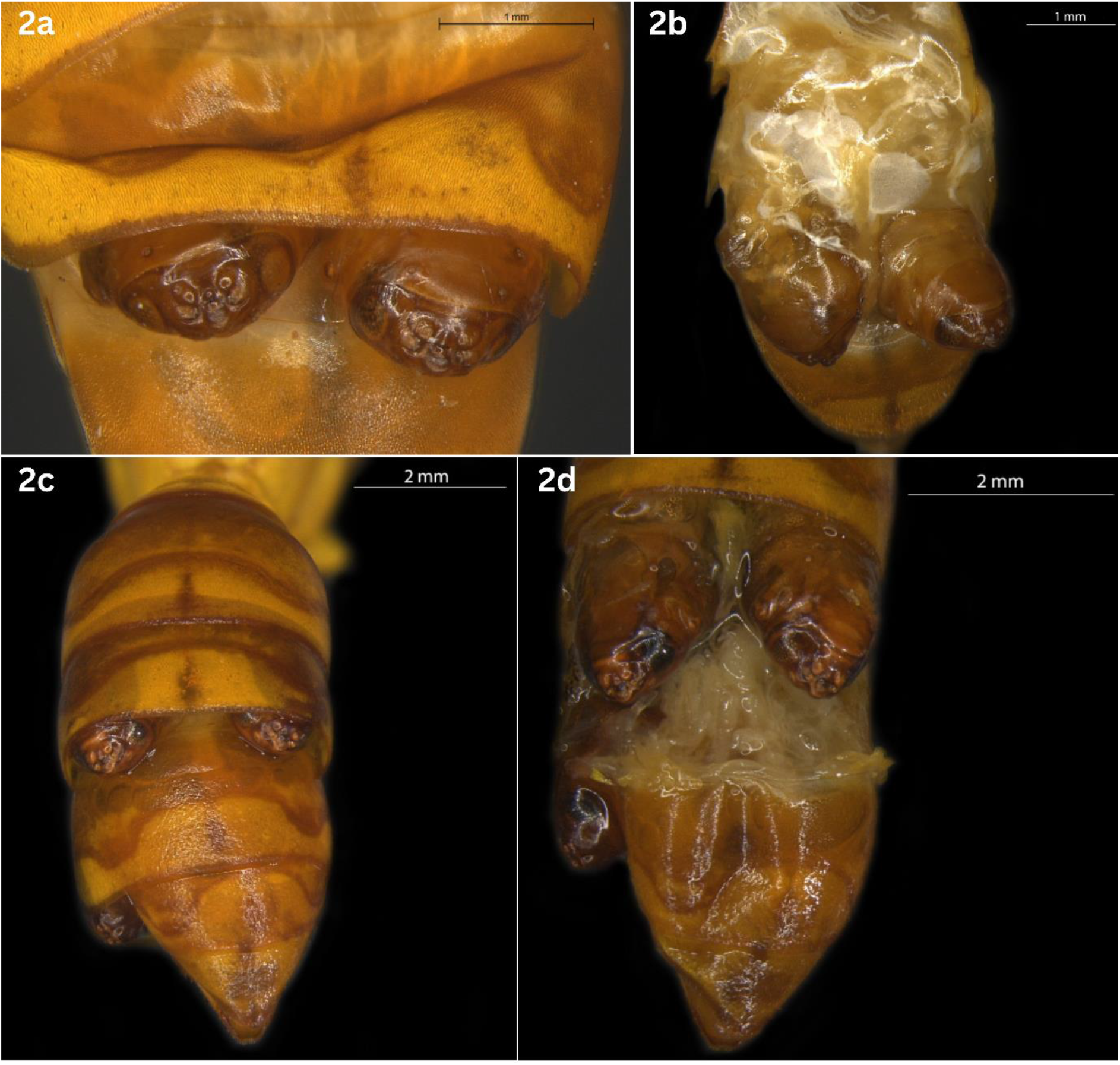
Abdomen of a female wasp with double infections (2a, b) and triple infections (2c, d). Left panel: Before sclerite removal, Right panel: After sclerite removal.

## Description

### Male puparium

The male *Xenos* puparium is elongated and oval shaped. The texture of the puparium is different in the anterior and the posterior side- it is darkly sclerotized and shiny anteriorly (till the thorax) and light brown and dull posteriorly (Fig 3). Length 4.53 ± 0.41 mm (n=7); Width 1.50 ± 0.13 mm (n=7). The anterior part is blunt and the posterior part bears the impression of the genitalia. Clear impressions of antennae, compound eyes (darkly sclerotized ommatidia) and mouthparts are visible through the puparium (Fig 3-4). The labial part is protruding below the distinct frontal, clypeal and labral region (Fig 4b).

**Figure 3.**
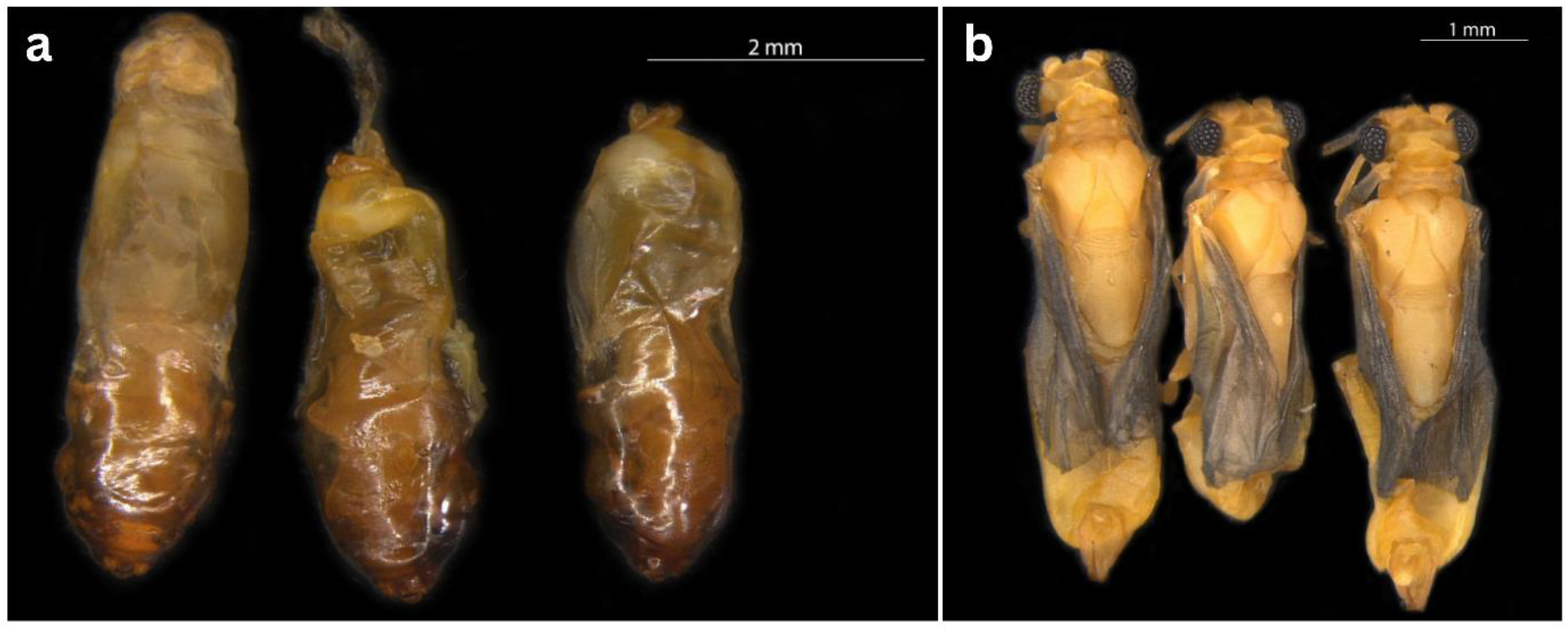
**a.** Puparia collected from a triple infected wasp (shown in Fig 2b). The anterior sides, represented by the dark brown, translucent cuticle are directed downward. 3b. Males dissected out of the puparia shown in 3a.

**Figure 4.**
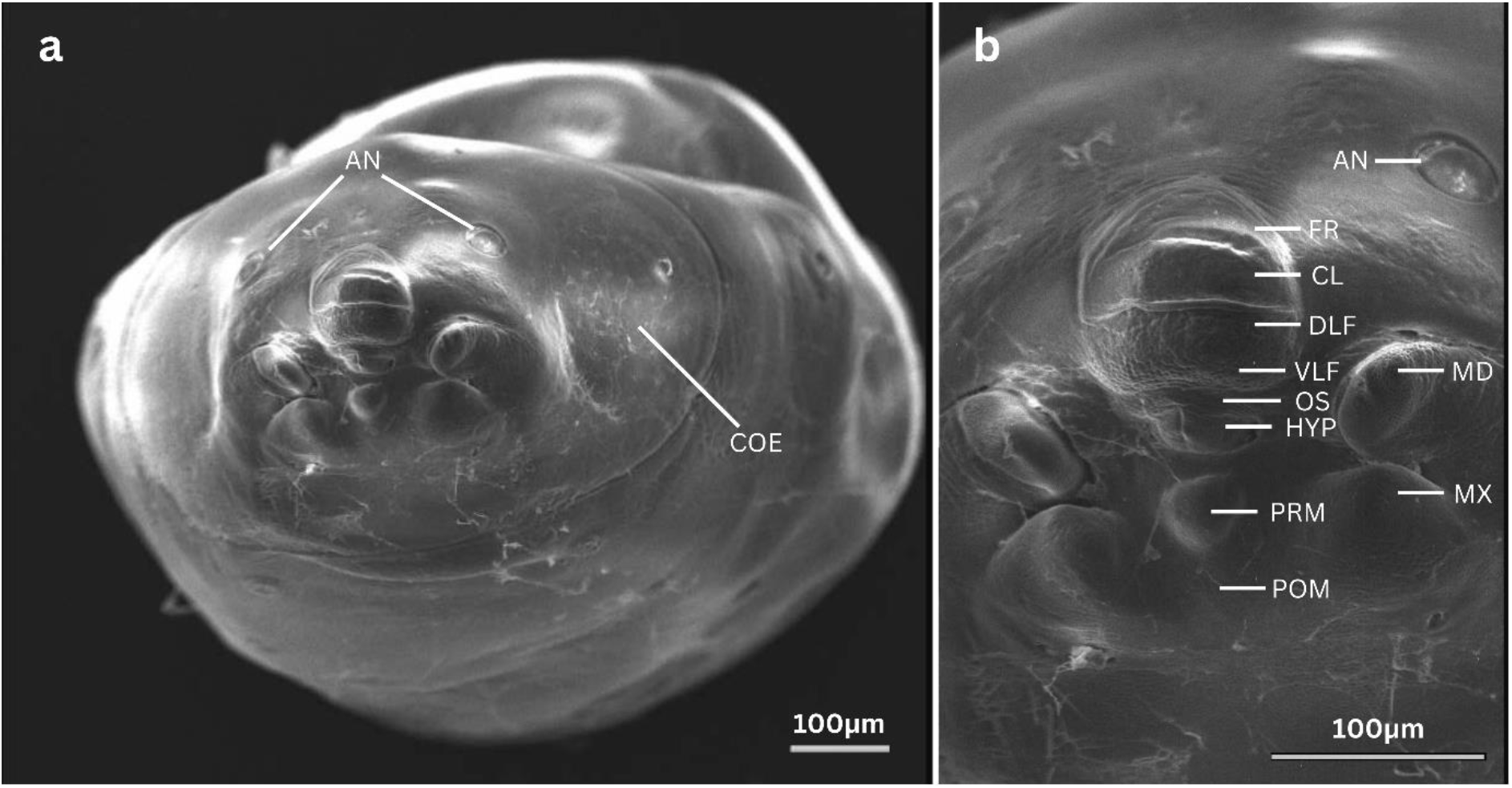
**a**.SEM image of the male puparium **b**. SEM image of the cephalotheca of the male puparium showing the impressions of the mouthparts and antenna. COE- Compound eye, AN- Antenna, FR- Frontal region, CL- Clypeus, DLF- Dorsal labral field of labral area, VLF- Ventral labral field of labral area, OS- Mouth opening, HYP- Hypopharynx, PRM- Prementum, POM- Postmentum, MD- Mandible, MX- Vestige of maxilla.

### Adult Males (Fig 5-6)

**Length:** Length varies according to the available space. In case of multiple infections, the males can be of different sizes (Fig 3). Anterior to posterior end: 4.12 ± 0.6 mm (n=7).

**Figure 5.**
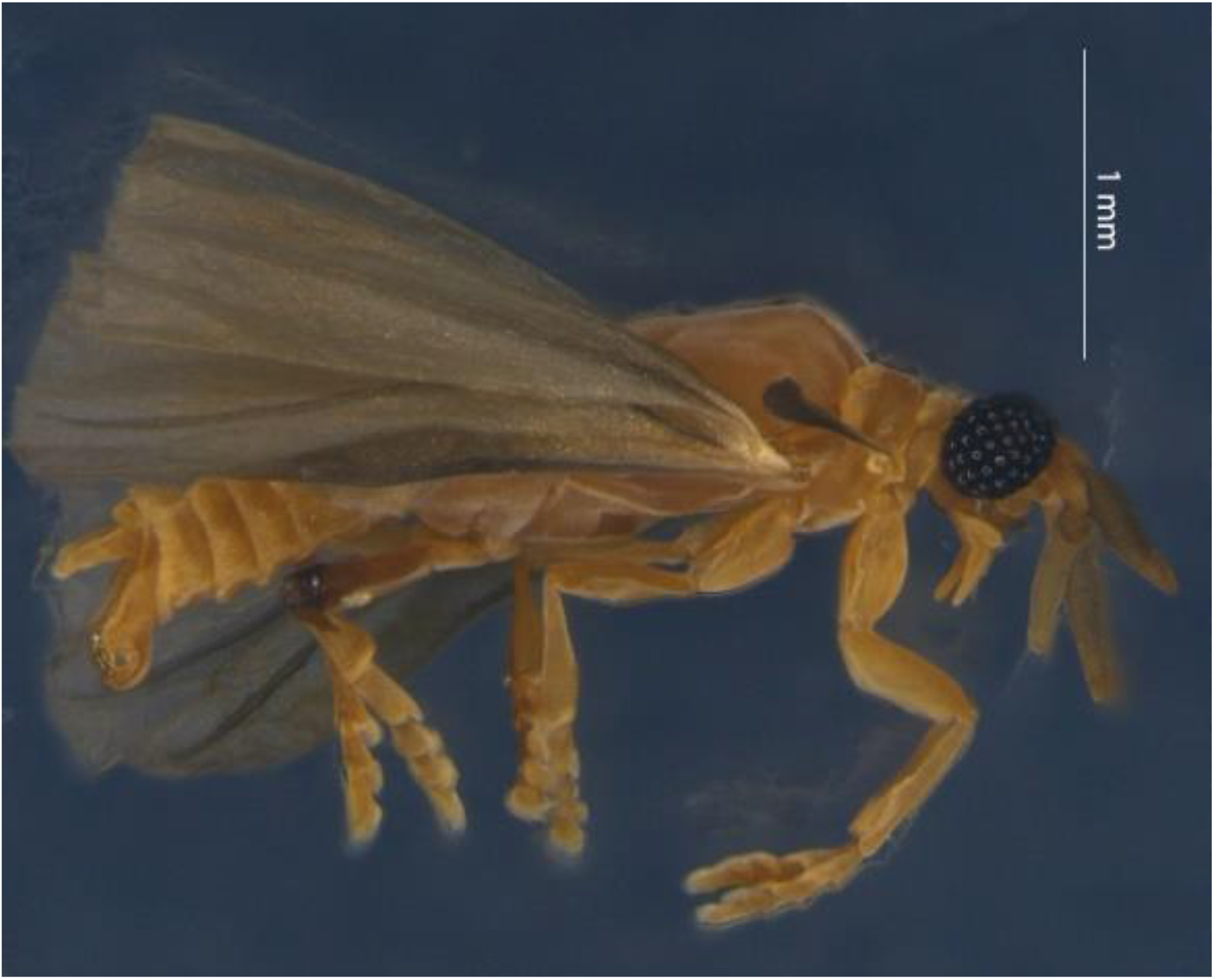
Male *X. gadagkari* eclosed naturally from the puparium.

**Figure 6.**
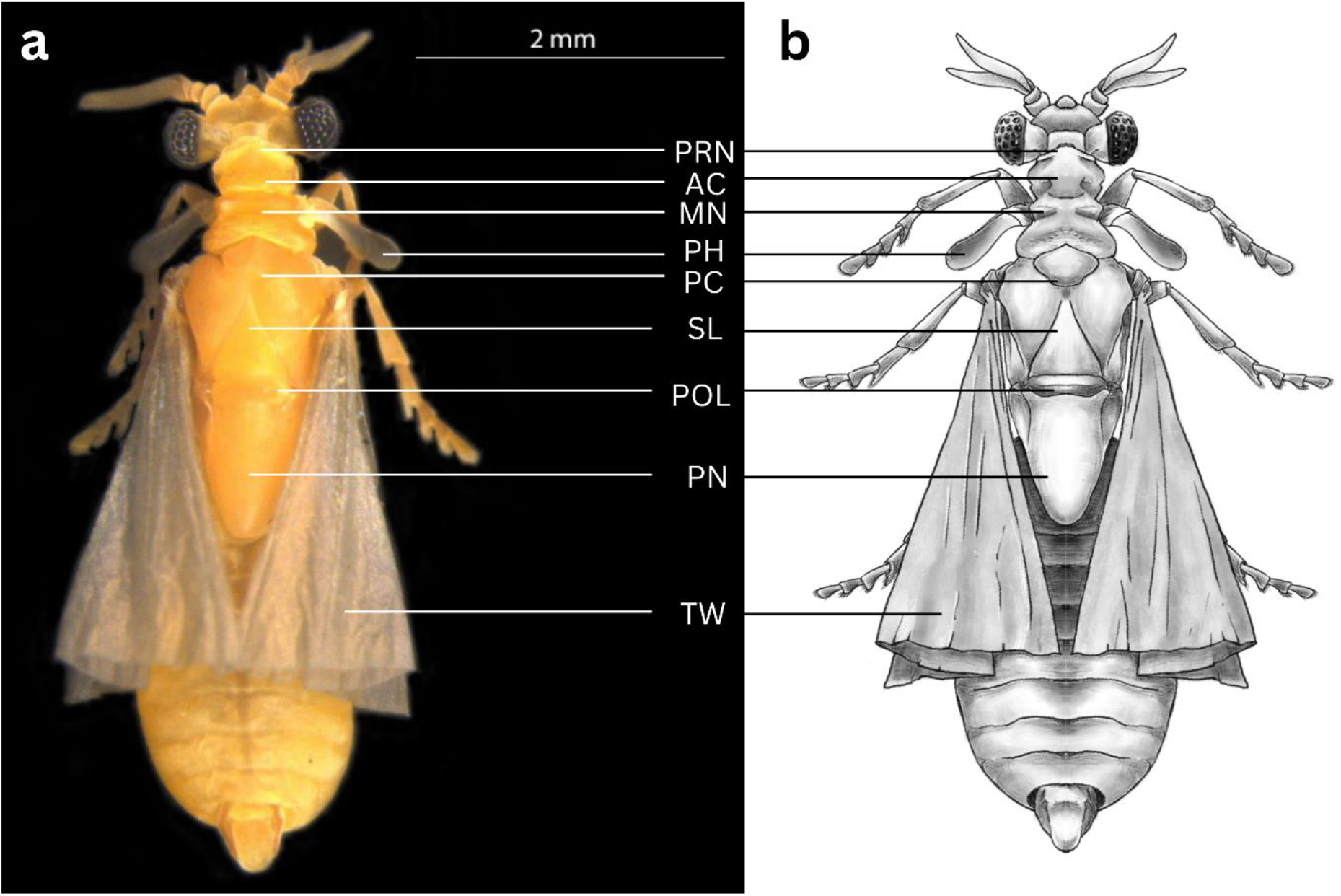
**a** Live male *X. gadagkari* taken out of puparium. **b.** Illustration of a male *X. gadagkari.* PRN- Pronotum, AC- Acrotergite, MN- Mesonotum, PC-Prescutum, SL-Scutellum, POL- Postlumbium, PN- Postnotum, PH- Pseusohalteres, TW- Twisted flying wing.

**Coloration:** Body and legs are bright yellow (honey gold) in colour (matching with the host *Polistes wattii’s* colour). Antennae are darker than the body, almost brownish. Halteres and metathoracic wings are semitransparent, greyish (lint) in colour (Fig 6).

**Head:** Hypognathus (Fig 6), wider than the first two thoracic segments. Head width including the compound eyes: 0.634mm ± 0.054 (n= 6)

**Compound Eye:** Raspberry like eyes with protruding ommatidia/eyelets. Approximately 65-70 (68 in left eye 66 in the right eye of one specimen) ommatidia are present in each eye. Ommatidia are distinctly outlined with tufts of small hair or microtrichia (Fig 7-8).

**Figure 7.**
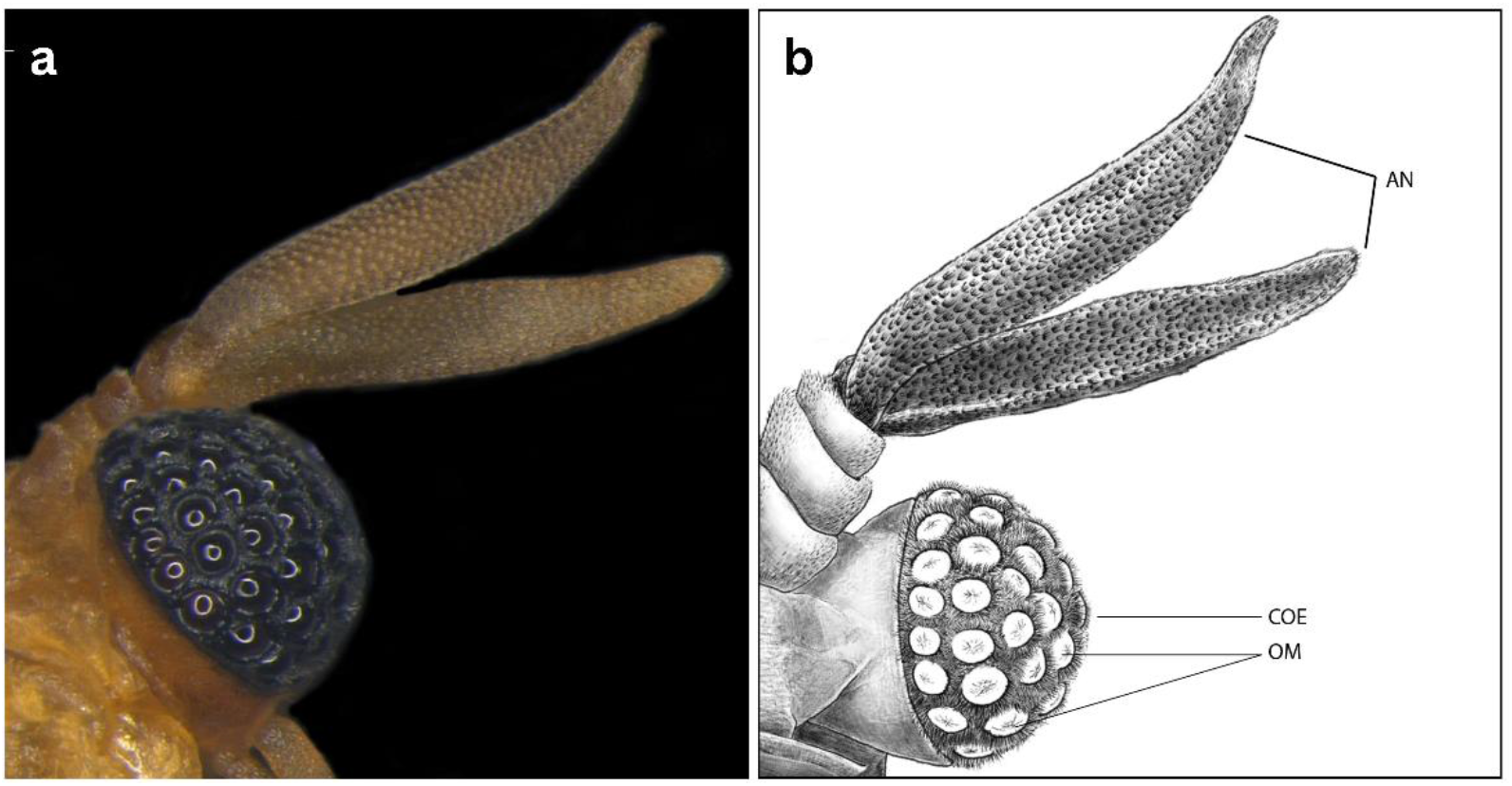
**a** and **b.** Compound eye and Flabellate antenna in male *X. gadagkari.* AN- Antenna, COE- Compound eye, OM- Ommatidia.

**Figure 8.**
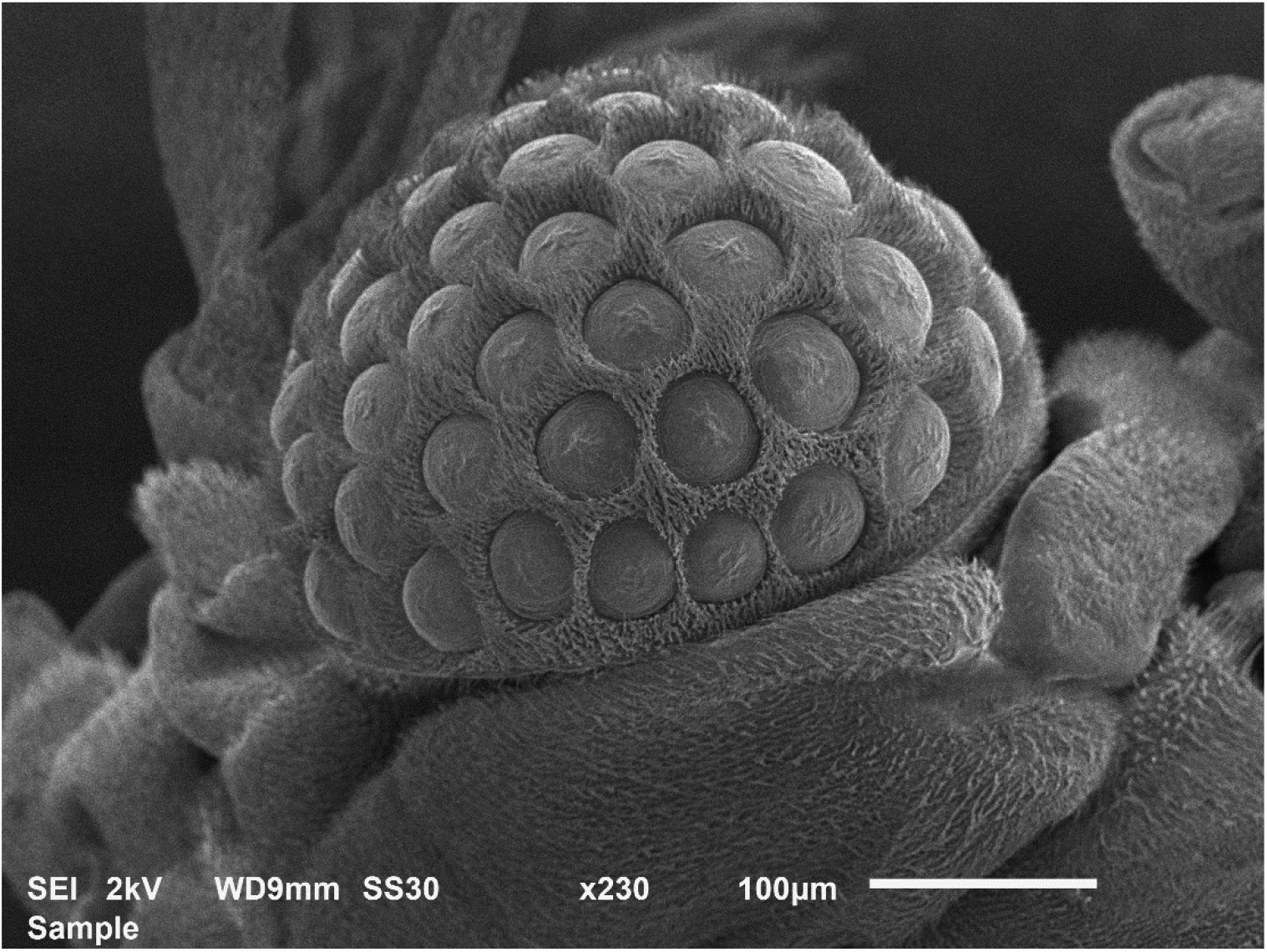
SEM image of the compound eye of male *X. gadagkari*.

**Antenna:** Flabellate antenna with four segments- scape, pedicel and two segmented flagella. Scape and pedicel are of equal width. Flagellar segments are almost equal in length and width. The whole antenna bears minute hairs (Fig 7, 9).

**Figure 9.**
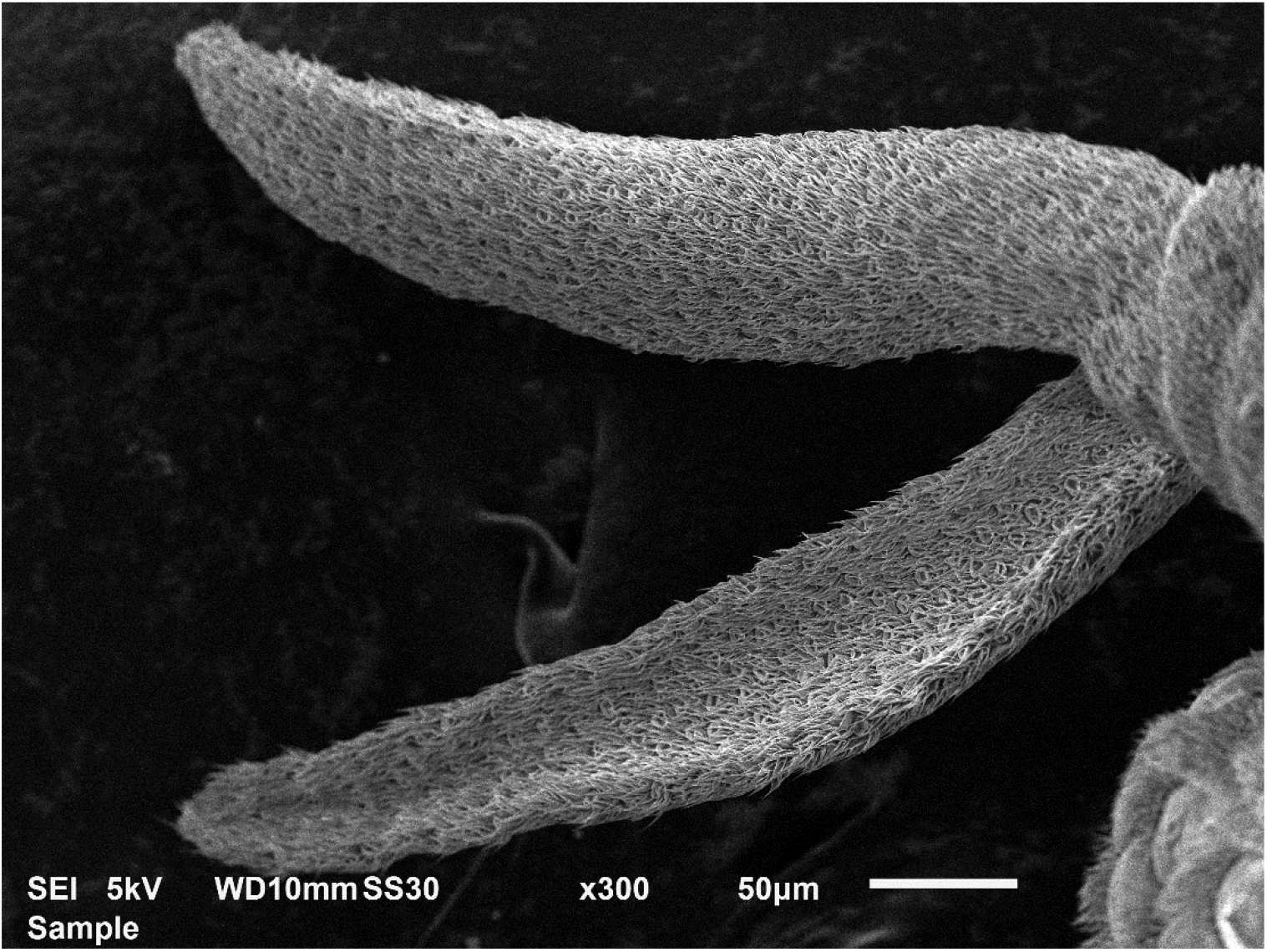
SEM image of the antenna of male *X. gadagkari*.

**Mouthparts:** A pair of mandibles and maxillae are visible. Mandibles are elongated and blade-like. Maxillae are two-segmented. The distal segment is darker than the proximal segment (Fig 10, 11)

**Figure 10.**
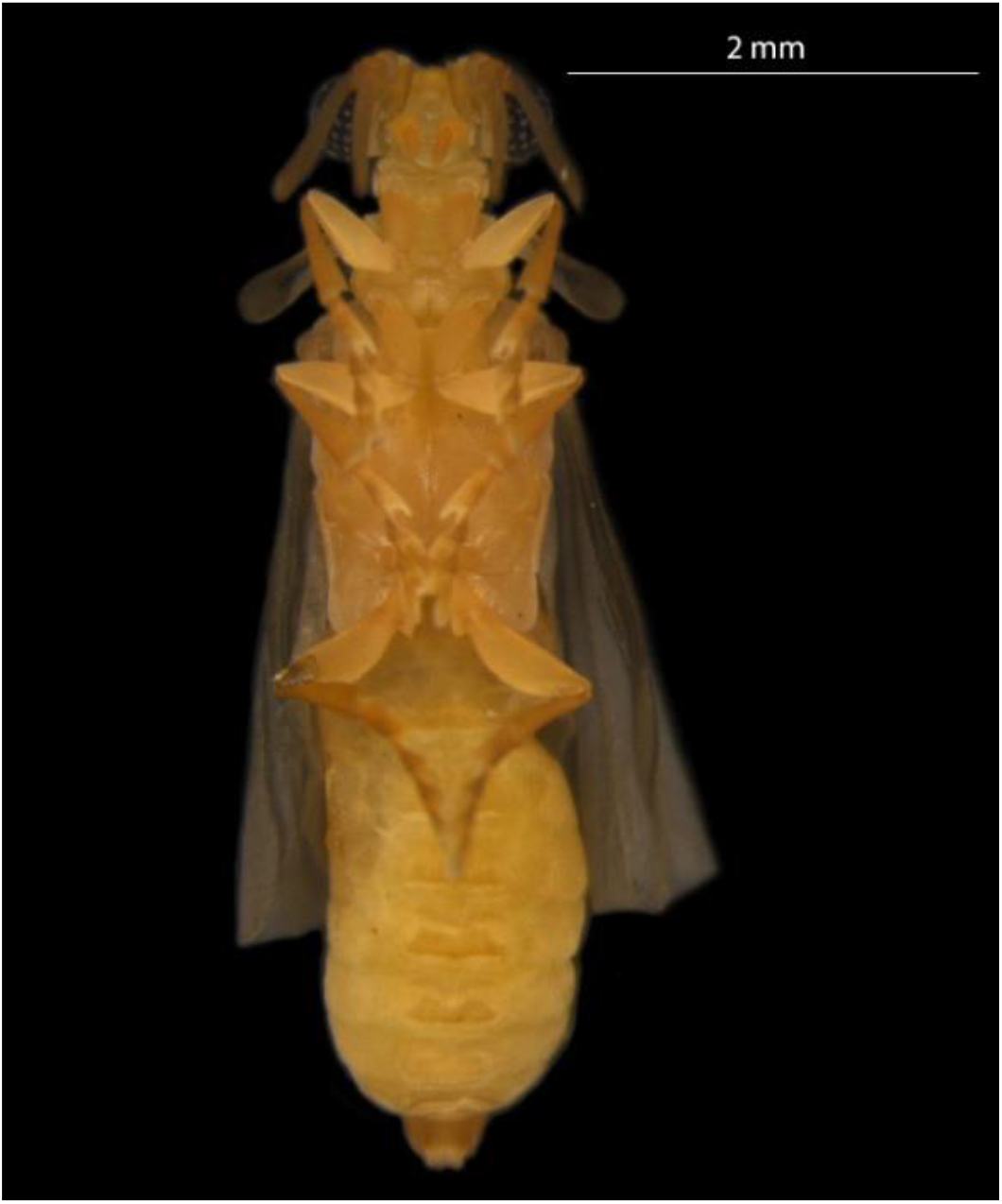
Ventral view of the male *X. gadagkari*.

**Figure 11.**
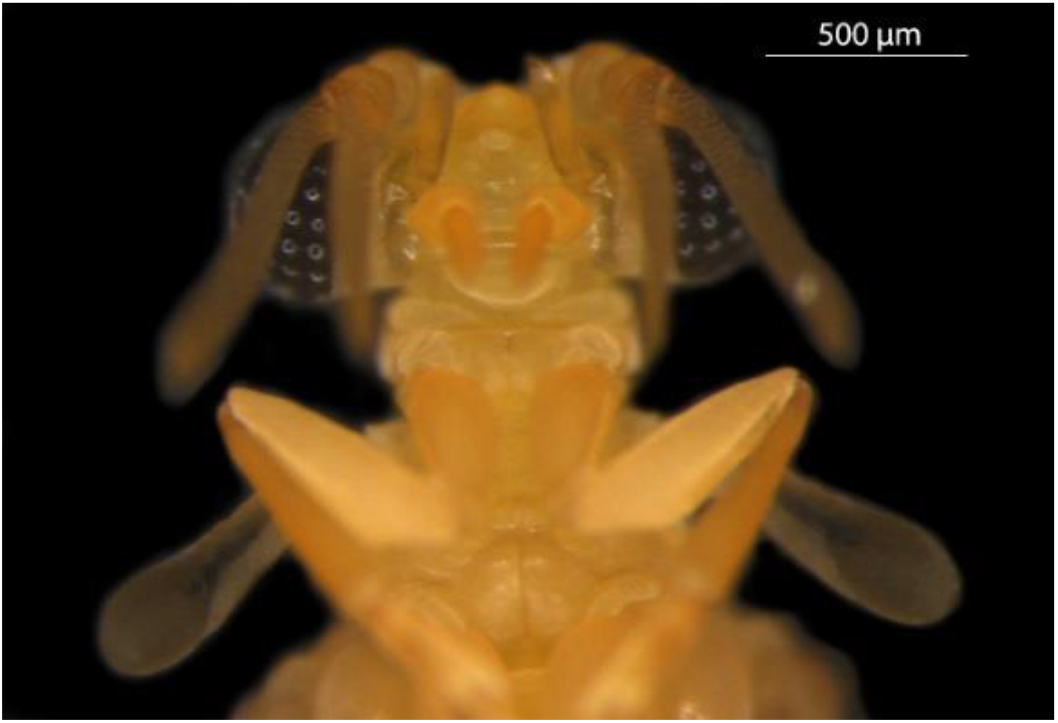
Mouth parts of male *X. gadagkari*.

**Figure 12.**
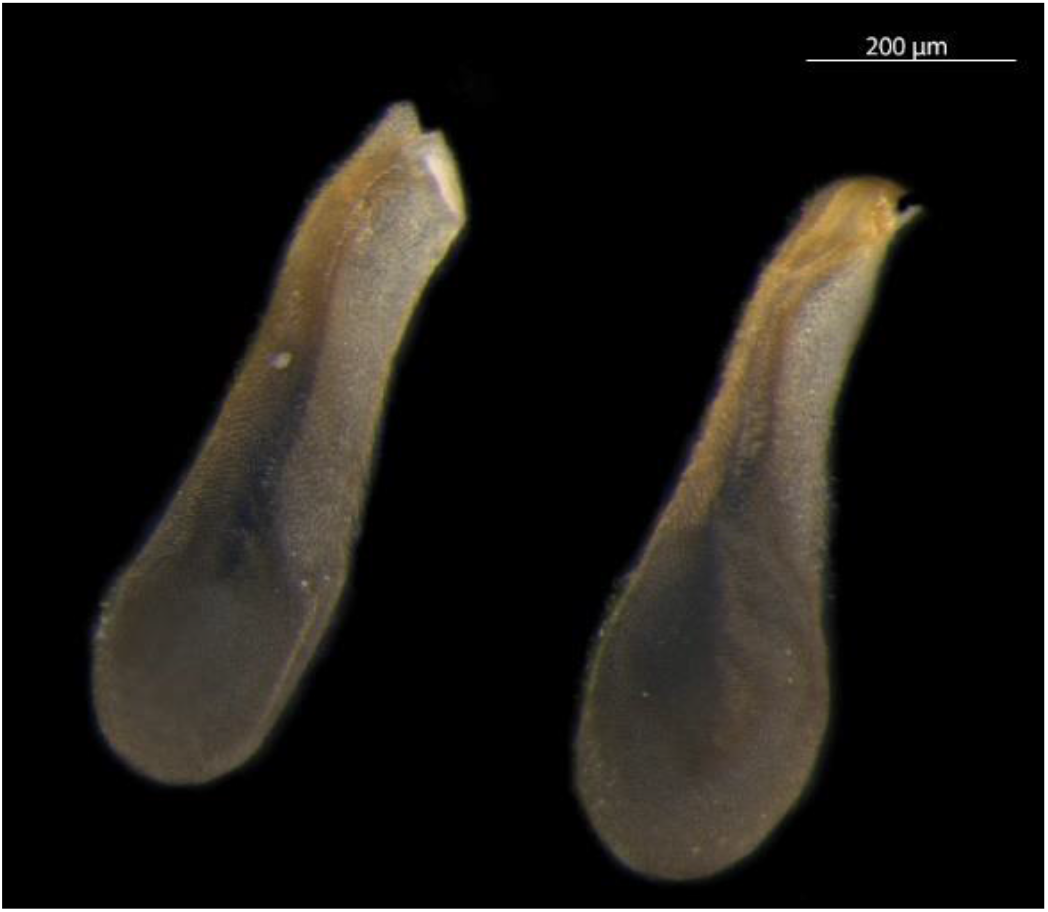
Pseudohalteres of male *X. gadagkari*.

**Prothorax:** Pronotum is trapezoid in shape, elevated dorsally; the average length of pronotum is 0.066mm ± 0.01 (n=6), swollen with lateral depressions (Fig 6).

**Mesothorax:** Mesonotum is visually wider than the pronotum and elevated dorsally (0.115mm ± 0.05, n=6. Fig 4). Forewings are modified into pseudo-halteres on both sides. Pseudohalteres are distally wide and bent both anterior and posteriorly into a boat-like structure (Fig 6).

**Metathorax:** Metathorax is more elaborate than the pro- and mesothorax. Metanotum length (1.523mm±0.054, n=6), Prescutum is pentagonal with a central longitudinal ridge and two depressions on both sides. Metathorax bears a pair of large fan-like wings (folded at rest) (Fig 6).

**Legs:** Fore, mid and hind legs are made of coxa, trochanter, femur, tibia and tarsus with four tarsomeres. Pretarsus is absent. All tarsomeres have hairy, oval, adhesive pads. The first tarsomeres of all legs have a pit (Fig 13).

**Figure 13.**
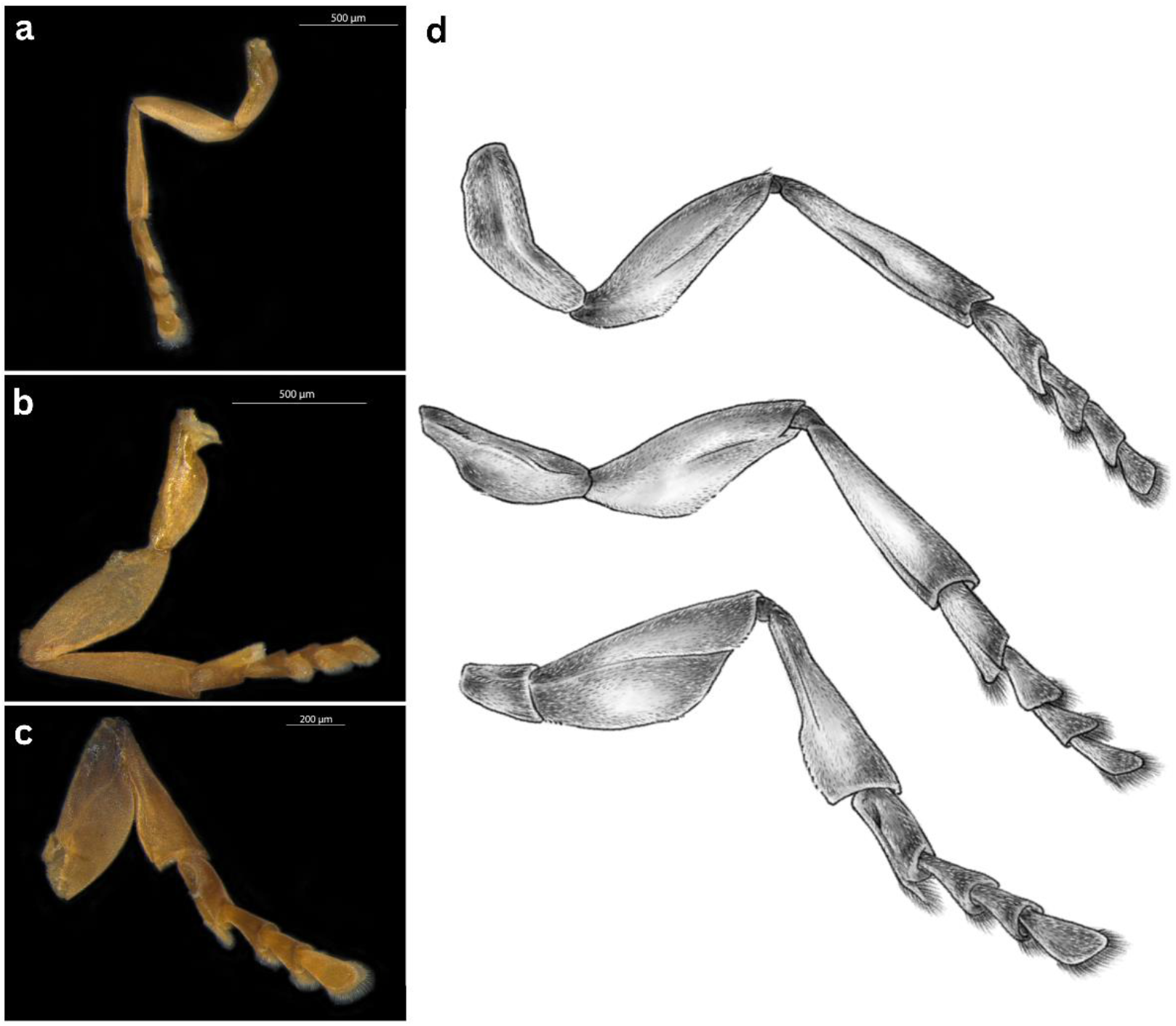
**a.** Proleg, **b.** midleg and **c.** hindleg of male *X. gadagkari.* **d.** Illustrations of the legs.

**Abdomen:** 8- segments are clearly visible, terminal segments are modified into genitalia (Fig 6).

### Neotenic Female

The female body size varies from host to host. Body length is 8.52mm ± 0.90 (n=6). Cephalothorax width is 1.38mm±0.13 (n=7), abdominal width is 2.22mm±0.23 (n=7). Colour: Cephalothorax dark brown, abdomen creamy white - light yellow with a central birthing canal with variable numbers of planidium larvae, which are clearly visible through the transparent cuticle of cephalothorax and abdomen, Fig 14-15). The birthing canal can be white or light brown before the hatching of the planidium larva but they appear darker brown with the increasing number of maturing larvae (Fig 14). The cephalothorax of the female is further described under the differences with *X. hebraei* and *X. vesparum*.

**Figure 14.**
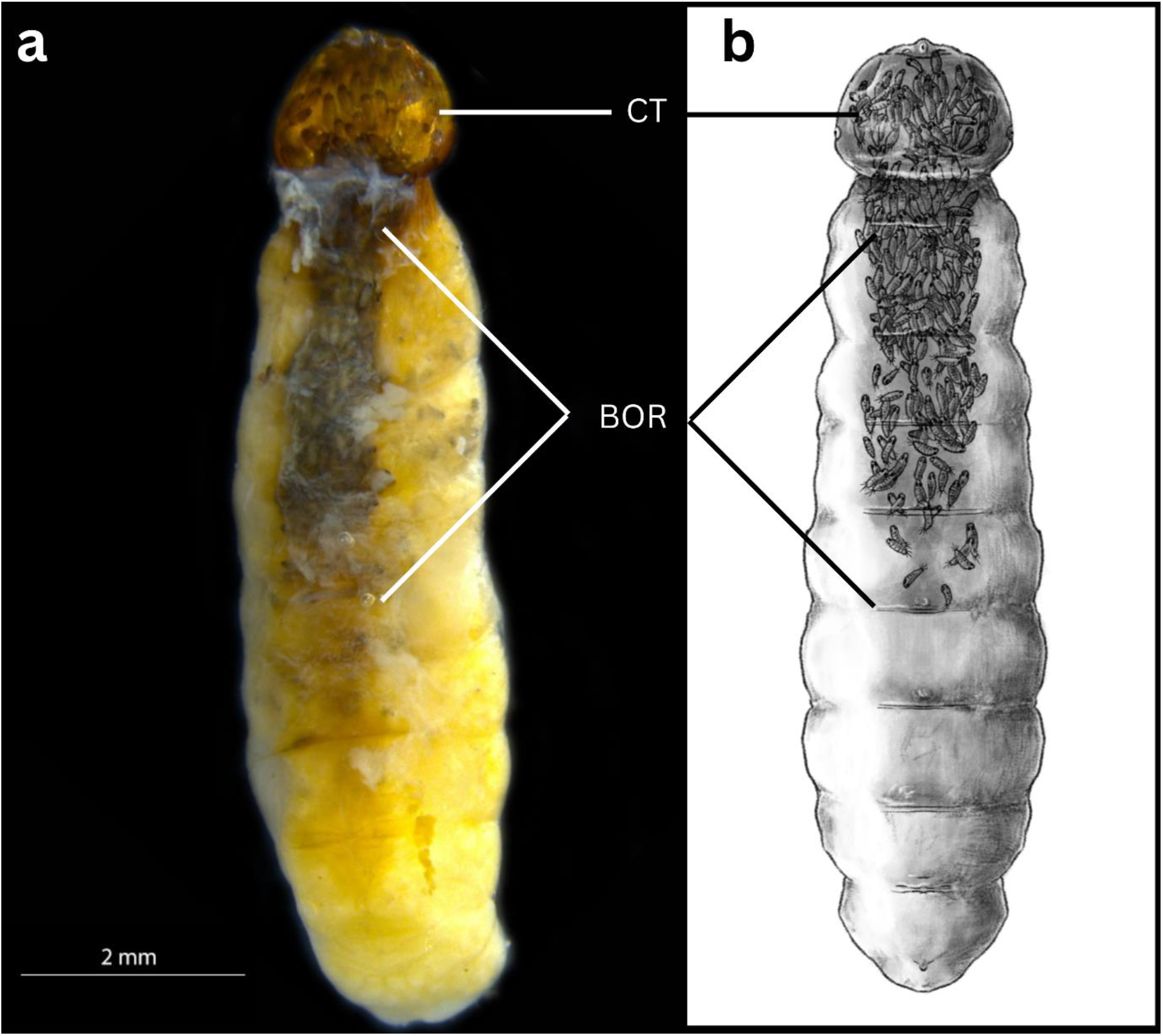
**a.** Fertilized female *X. gadagkari* with planidium larvae collected from *P. wattii.* **b.** Illustration of female *X . gadagkari.* CT- Cephalothorax, BOR- Birthing organs.

**Figure 15.**
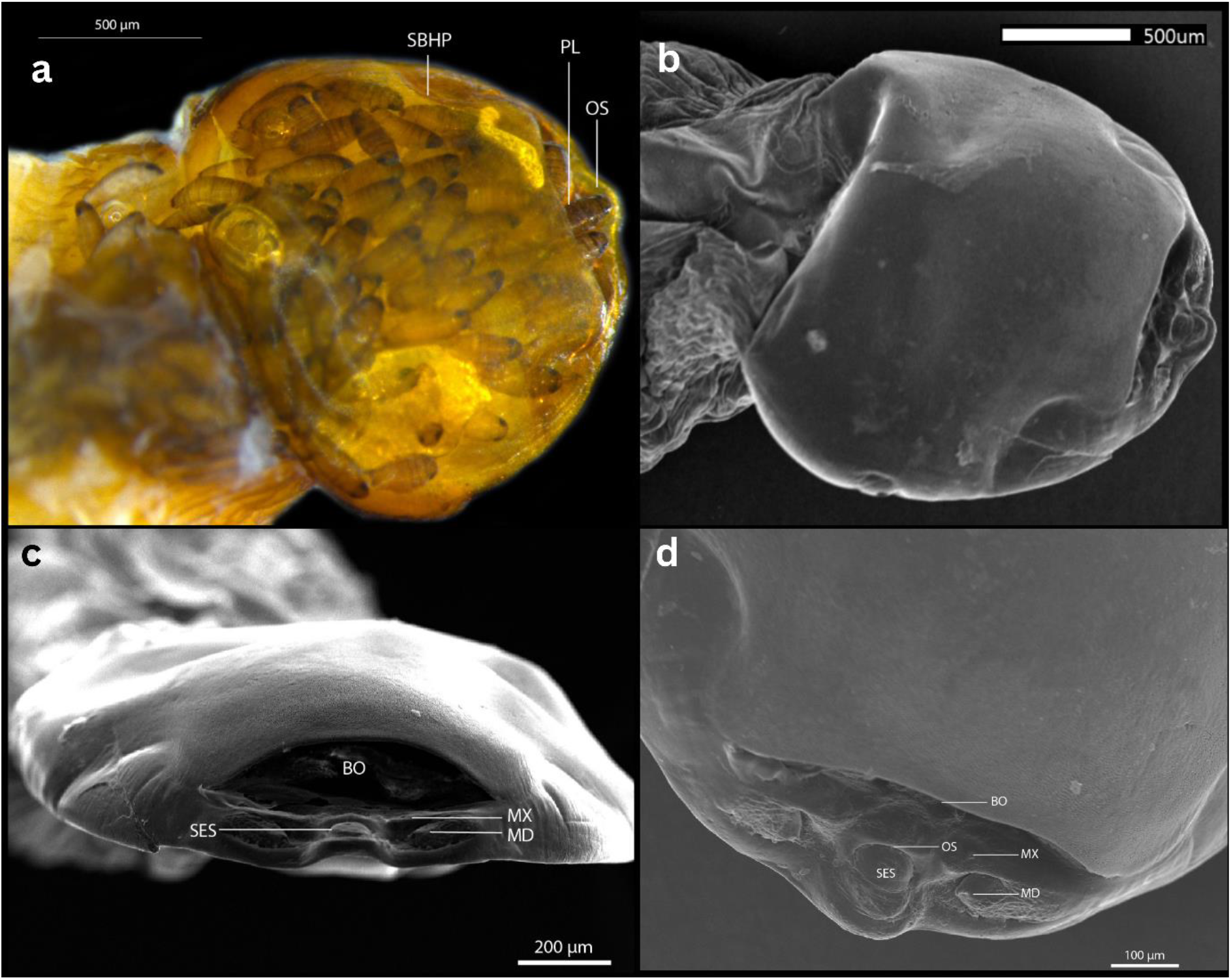
**a.** Cephalothorax of female *X. gadagkari* with planidium larvae collected from *P. wattii.* **b, c** and **d** SEM images of the female cephalothorax. SBHP- Segmental border between head and prothorax, PL- Planidium larvae, OS- Mouth Opening BO- Birth opening, SES- Semicircular field possible of labral region, MX- Maxilla, MD- Mandible.

### Planidium Larva

Varied numbers of first instar planidium larvae (also known as triangulin larvae) were found inside the neotenic female (Fig16a). Planidium larvae are approximately 200µm in length from anterior to posterior end. They are characterized by sclerotized exoskeleton, two black eyespots and two pairs of caudal setae (Fig 16b).

**Figure 16.**
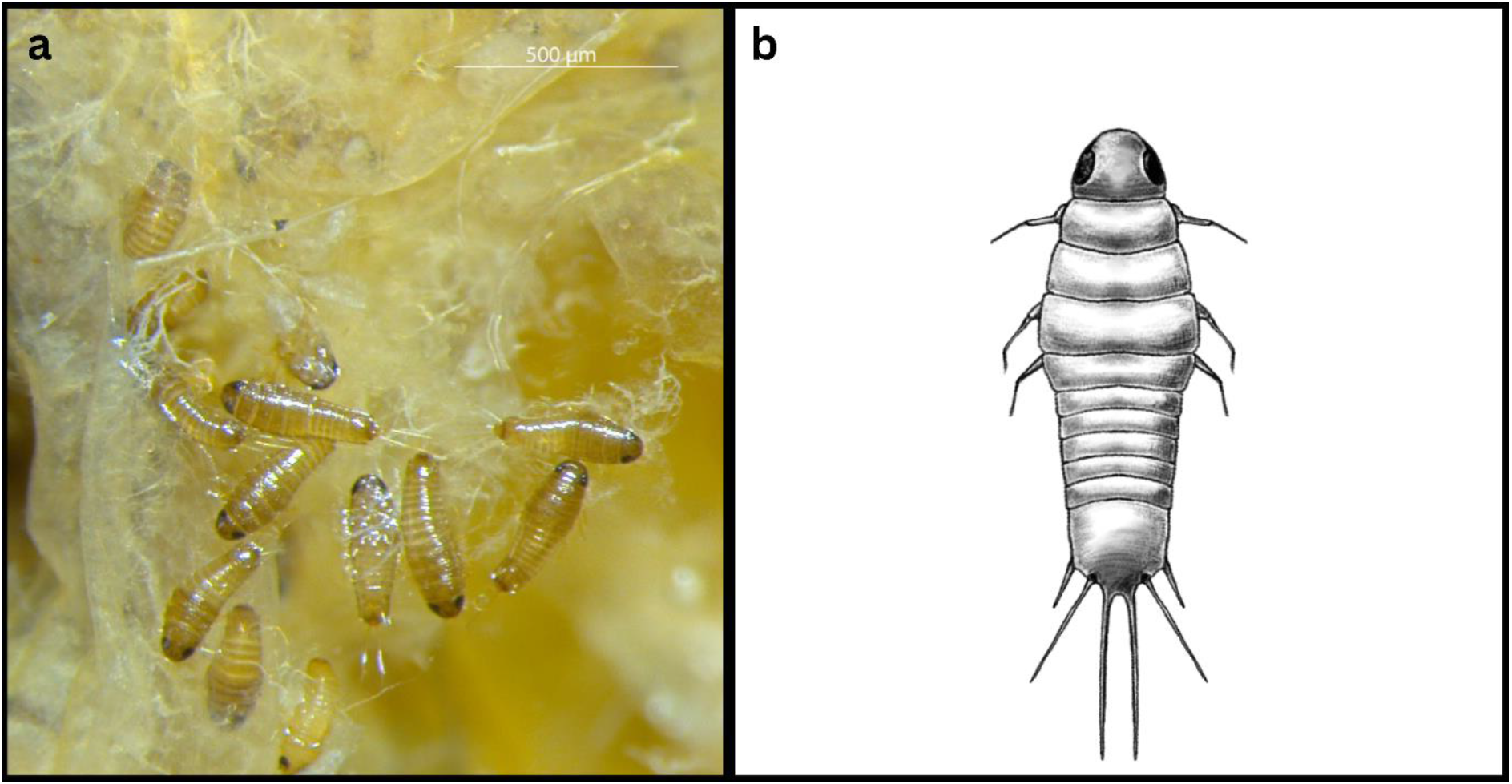
a. Planidium larvae inside the cephalothorax and exiting through the gonopore of female *X. gadagkari.* b. Illstration of a single planidium larva.

## Taxonomy

**Xenidae Saunders 1872**

***Xenos Rossius*, 1793**

***Xenos gadagkari*, Sen and Nain sp.nov.**

lsid:zoobank.org:pub:A1D78367-F2EB-4408-A077-5EA8888D6B60

### Phylogenetic analysis

The phylogenetic analysis of *X. gadagkari* was done with several partial fragments of the mitochondrial COI gene sequences of other species obtained from NCBI. *X. gadagkari* forms a clade with *X. vesparum* among several other strepsipterans from different Polistine wasps (Fig 17). The clade is also supported by high posterior probabilities (100) indicating strong reliability of the clade. The *X*. *gadagkari* sequences show a synonymous site divergence of 79% (with 68% divergence across all the sites considered) from *X. vesparum*, indicating a very different evolutionary history.

**Figure 17.**
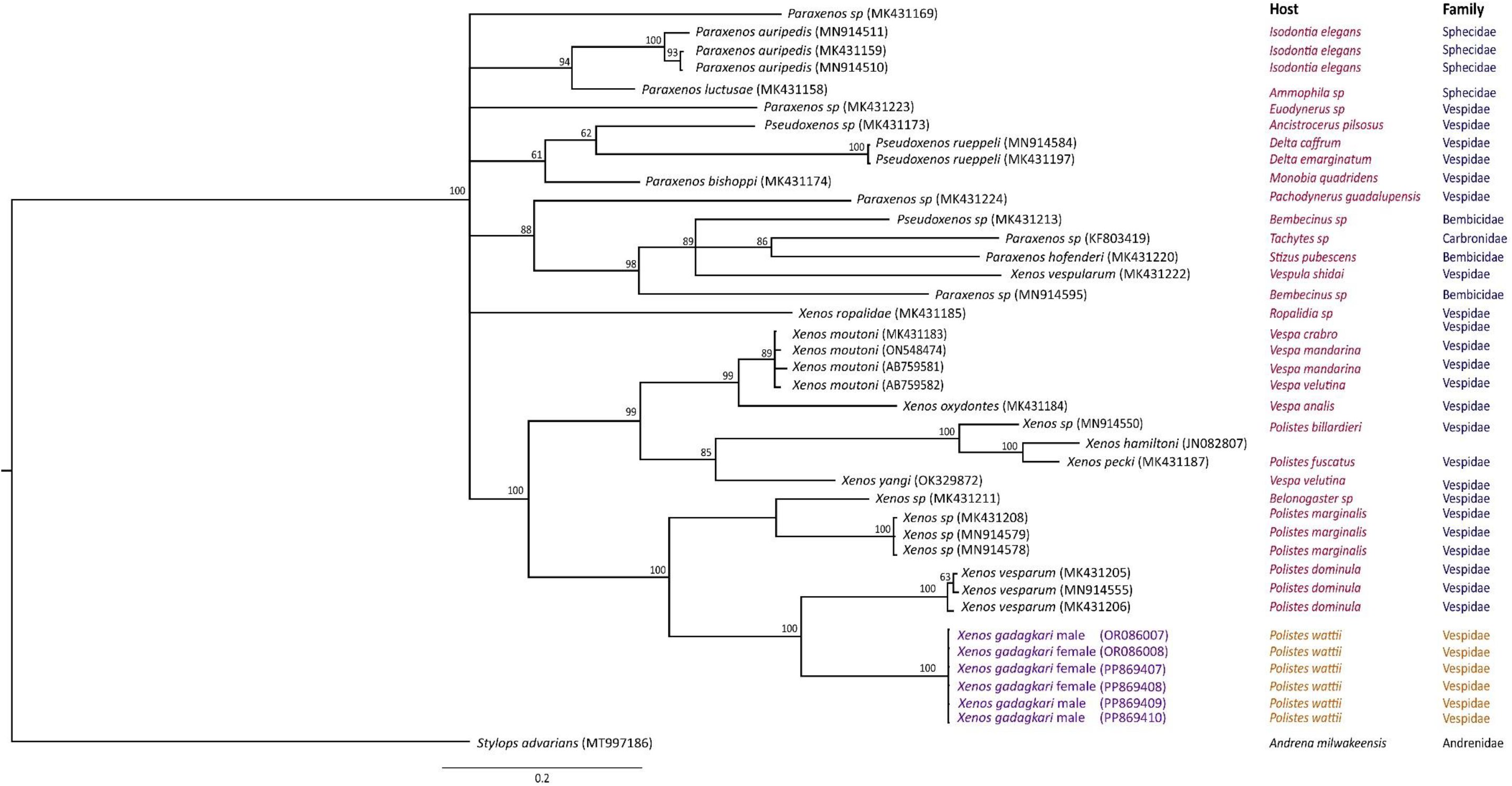
A Baysian phylogenetic tree of partial mitochondrial *CO1* sequences constructed using GTR+g_i model in MrBayes for different species of Xenidae family. *X. gadagkari* sequences generated for this study are represented in purple. Homologue from *Stylops advarians* (host: *Andrena milawaukeensis*) was used as an outgroup. Numbers above the nodes indicate clade credibility values. Names of the hosts and their family are given against the sequences. Host name was not available for *X. hamiltoni* in the paper associated with the accession number. However, *X. hamiltoni* is usually found in *Polistes carnifex* (Kathirithamby & Hughes, 2006).

### Differences with X. hebraei and X. vesparum

*X. gadagkari* is phylogenetically closest to *X. vesparum* (according to currently available sequences). However, *X gadagkari* appears to be visibly different from *X. vesparum* particularly due to the drastic difference in colouration. In the female cephalotheca, the mandibles of *X. gadagkari* are positioned in a depression larger than their length, while in *X. vesparum* the depression around the mandibles is smaller than the length of the mandibles. Also, the lower margin below the semicircular field possible of labral region (SES) is elevated (Fig 15c) in *X. gadagkari,* while it is smooth in *X. vesparum* (Fig 2D in Richter et al 2017)

As the description of the adult male *X. hebraei* is unavailable, it is impossible to compare it. However, according to the description of the female cephalothorax of *X. hebraei* (Kinzelbach 1978) ‘the clypeo- labral region protrudes less than in any other of the species of western Palearctic’. The protrusion of the clypeo-labral region in *X. gadagkari* is more pronounced than *X. vesparum* (Fig 15d of this study compared with Fig 2D of the study on *X. vesparum* (Richter et al., 2017). Kinzelbach (1978) further described that ‘the posterior delimitation of the brood aperture is shifted far forward’ while the lateral margins of the brood aperture reach the anterior margin of the head. In *X. gadagkari,* neither of these two characters is different from *X. vesparum* i.e. the posterior delimitation of the brood aperture is not shifted forwards and the lateral margins of the brood aperture end way before reaching the anterior margin of the head. Although body size may vary from specimen to specimen, a morphometric and colour comparison of *X. gadagkari* with the available data from *X. vesparum* and *X hebraei* is compiled in table 2.

**Table 2.**

|  | *X. gadagkari*<br>(this study) | *X. hebraei*<br>(Kinzelbach 1978) | *X. vesparum*<br>(Beani et al 2021) |
| --- | --- | --- | --- |
| Male Body length | 4.12±0.6 mm (n=7) | 5 | Not reported |
| Male Puparium length | 4.53±0.41 mm (n=7) | 6.5 mm | 4.9±0.03 mm (n=7) |
| Male puparium width | 1.50±0.13 mm | 1.5 mm | 1.66±0.03 mm (n=7) |
| #Ommatidia | 65-70 | 80-82 | 65 |
| Male body colour | Yellow | - | Dark brown to black |
| Female cephalothorax width | 1.38±0.13 (n=7) | 2.0 mm | 2.48±0.05 mm (n=34) |
| Female body length | 8.52±0.9 (n=6) | 7.5 mm | 7.93±0.12 mm (n=34) |
| Female body colour | Creamy white-light yellow with a dark brown birthing canal | Yellowish Brown, | Creamy white with brown birthing canal |
| Cephalothorax colour | Dark brown |  | Horny yellow to black-brown |

## Discussion

Lefroy (1909) reported the presence of *Xenos* in *P. hebraeus* (as mentioned by Green, 1912) from India but the species was not described. In a compilation of a preliminary checklist of species from Family Xenidae, Benda *et al* (2022) reported the presence of *X. hebraei* in *P. wattii* from Oman assuming the occurrence of the same species as present in *P. olivaceous* reported from Iraq (Kinzelbach; 1978). Kinzelbach’s study documented a short description of the female but lacked a description of the mature adult male. Therefore, there is no convenient way of knowing whether it is the same species as found by Kinzelbach or not. Our attempt to track the original holotype of *X. hebraei* was unsuccessful. The description of the female cephalothorax and the morphometric description of the female of *X. hebraei,* differ from the species found in *P. wattii.* Therefore, it appears fit to report *X. gadagkari* as a new species with a full description of male puparium, adult male, neotenic female and planidium larva with images and illustrations and a phylogenetic analysis with other available *Xenos* sequences.

*X. gadagkari* is commonly found in the aggregations of *P. wattii* wasps in May and in wasps in the late colony phase. The parasite infects both sexes of the host but the role of male hosts in the spread of the parasite is not known. Here we confirm the presence and abundance of *Xenos* in *P. wattii* in India and it would be interesting to study the biology of *Xenos* and its effect on the nesting biology, behaviour and morpho-physiology of an Indian *Polistes.* Since the behavioural, physiological or molecular aspects of *Xenos-Polistes* association have not been studied in India, *P. wattii - X. gadagkari* association can be a useful model for such investigations. The holotypes of *X. gadagkari* will be deposited at the entomological collection of the Zoological Survey of India, Kolkata, India.

## Acknowledgement

The authors thank Prof Jeyaraney Kathirithamby for helpful discussions. The authors are thankful to the Department of Forests and Wildlife, Chandigarh Administration and the Department of Forests and Wildlife Preservation, Punjab, for the permit and NOC to collect wasps. The authors also thank Atharva Deshpande for making the illustrations used in this paper and Rajbir Kaur, Arpita Nath and Kunika Malhotra for helping with the initial observations on this system.

## Author contribution

DN collected and dissected the wasps and measured the *Xenos* specimens, DN, AR and RR carried out the sequencing and phylogenetic analysis. RS discovered the species, designed the study, supervised data collection and analysis and wrote the paper with DN. All authors have read and contributed to the final edit of the manuscript.

